# Selection for propagule formation in prebiotic chemical ecosystems

**DOI:** 10.64898/2026.09.08.750225

**Authors:** Nicole Anderson, David Baum

## Abstract

Biological individuals on Earth today are cellular, but the simplest cells are too complex to have arisen spontaneously. We explore the hypothesis that the emergence of biological individuality might have been driven by selection for different molecules to move through space together by forming a propagule. We predicted that propagule formation will be advantageous to an autocatalytic system primarily when 1) sites supporting autocatalysis are distantly spaced and 2) the environment undergoes periodic disturbances. We tested this hypothesis using computational reaction-diffusion models of surface-catalyzed autocatalytic systems in patchy, disturbance-prone environments. We considered two competing autocatalytic cycles, each composed of mutualistic subcycles, one of which diverts flux to the production of propagules. Our analysis shows that propagules tend to be detrimental in well-mixed environments or environments where viable surfaces are close enough together to be reliably seeded through single-species dispersal. However, propagule formation is advantageous when long stretches of empty space must be crossed to reach new resources and/or when environmental disturbances are frequent. These findings suggest spatial structure can selectively favor reliable forms of codispersal. Evolution in response to this mechanism could potentially explain the emergence of vesicles that transported many autocatalytic species and eventually became independent protocells.

**Author summary:** Cells on Earth today are too complex to have emerged spontaneously, meaning they must have developed through a gradual evolutionary process. Early evolution could have occurred through selection on self-amplifying sequences of chemical reactions called autocatalytic cycles. These cycles resemble processes in modern life, such as metabolic cycles and DNA replication. Understanding how these chemical cycles became bound into cells is vital for understanding the origin of life from prebiotic chemistry. We suggest that selection for codispersing between favorable sites by forming a propagule could have eventually led to the emergence of cells. We use computer models to test the idea that propagule formation is favored when the distance between resource-rich sites is long and the environment is periodically disturbed. By showing that realistic environmental settings drive the emergence of propagules, which could then serve as targets of selection for traits such as growth and division, our research helps explain the emergence of the first biological individuals.

## Introduction

The origin of life is often envisioned as a process by which networks of chemical reactions become more elaborate until they develop lifelike qualities, often defined as the ability to sustain themselves and undergo Darwinian evolution (per the “NASA definition” of life [1]). On Earth, this process resulted in genetic mechanisms of inheritance and membrane-encapsulated cells, which are common to all known lifeforms. However, these systems are too complex to have arisen spontaneously, suggesting they must have arisen gradually through pre-genetic and/or pre-cellular evolution.

Non-genetic evolution involves transitions among alternative compositional steady states [2] in networks that include cycles of reactions that collectively increase the stoichiometry of a set of member species, known as autocatalytic cycles (ACs) [3]. Because ACs are self-maintaining as long as they are provided with resources, the activation of an AC in a system is heritable in the sense that it causes a potentially irreversible change in the state of the system [4, 5]. Therefore, while ACs alone do not have the capacity to evolve, sets of ACs contain information that can be acted upon by selection or change randomly with drift [6]. For selection to work, these sets must be spatially distinct from each other. Some models, such as GARD, solve this problem by assuming that ACs or sets of mutual catalysts start out bound into self-organized lipid assemblies capable of growth and division, meaning individuation arises spontaneously [7]. Others suggest that the environment imposes either a continuous or patchy spatial structure, such as distinct mineral surfaces or pores [8, 9]. Each patch or pore can then be treated as an “individual” with a compositional “genome” composed of its active ACs, with variable fitness depending on, for instance, the resistance of those ACs to environmental perturbation or the speed at which they spread to acquire new resources.

If we allow that prebiotic evolution can begin with individuals whose boundaries are defined by the external environment, we must explain how these environmentally separated units could become mobile individuals with self-maintained boundaries, fulfilling the definition of autopoiesis [10]. It has been hypothesized [11] that primordial selection could drive the emergence of propagules - structures that enable seeds from a set of interdependent ACs to move from one environmental patch to another. Over time, selection for increasingly reliable propagules would result in structures with self-maintained boundaries and the ability to divide; these would be protocells and predecessors to biological individuals. Propagules themselves might meet definitions of individuality based on the degree of codispersal between replicating molecules [12]. However, selection on propagule fitness would likely drive the evolution of additional attributes such as the cohesion and integration of their component parts, satisfying a variety of competing definitions as well [13].

A central driver of the emergence of propagules in this model is the selection for a tendency for different types of molecules to preferentially move through space together, or codispersal. Association between molecules might be achieved by containment into mobile propagules like vesicles, micelles, or coacervates. An even simpler form of propagule would be a large molecule or polymer formed by covalent bonds between the members of different ACs. Propagules should be favored especially when an AC requires multiple types of seeds to establish itself in a new location; in other words, when the cycle requires a composite seed [14]. One type of cycle which would require a composite seed would be one that is composed of multiple obligately mutualistic subcycles, each of which needs to be active for any to persist. Such networks of interdependent subcycles might not be capable of reaching resource-rich patches if they are distant enough that the probability of the seeds of all the subcycles arriving independently at the same time is low.

Here, we construct in-silico models of ACs to examine the spatial conditions and AC structures that favor propagule-forming cycles. We tested the prediction that propagule formation would be more advantageous when spatial and environmental constraints are more stringent, that is when sites capable of supporting autocatalysis are more distantly spaced and when the environment undergoes more frequent disturbances which clear sites of their autocatalytic species. We tested this prediction by analyzing reaction-diffusion models of competing ACs, one of which forms propagules while the other does not. As predicted, long distances between patches of resources and frequent environmental disturbances make propagule-forming systems more likely to persist and reach high concentrations. These findings demonstrate that the existence of cooperating chemical processes in spatio-temporally structured environments can help drive the emergence of individuals.

## Materials and methods

### Obligately mutualistic ACs and propagules

We consider ACs composed of sets of obligately mutualistic subcycles (Fig 1). Each AC contains a member species that can react with a generic food species, *F*, to produce a waste product and an intermediate. Each intermediate can react with the waste product of one of the other subcycles to produce two of the original member species. For simplicity, we assume that the food, intermediate species, and member species have the same mass, which ensures mass conservation. All reactions are reversible (e.g., two member molecules can react to generate one intermediate and the waste of another cycle). In such a network, member species can produce intermediates and waste so long as *F* is available, but the population of the member species will not increase unless the other subcycles are also present and active. We refer to the smaller autocatalytic rings depicted in (Fig 1) as subcycles, and to the entire network of mutualistic subcycles as an AC, or simply a cycle. The set of all possible reactions in the world is referred to as a chemical reaction network or CRN.

**Fig 1.**
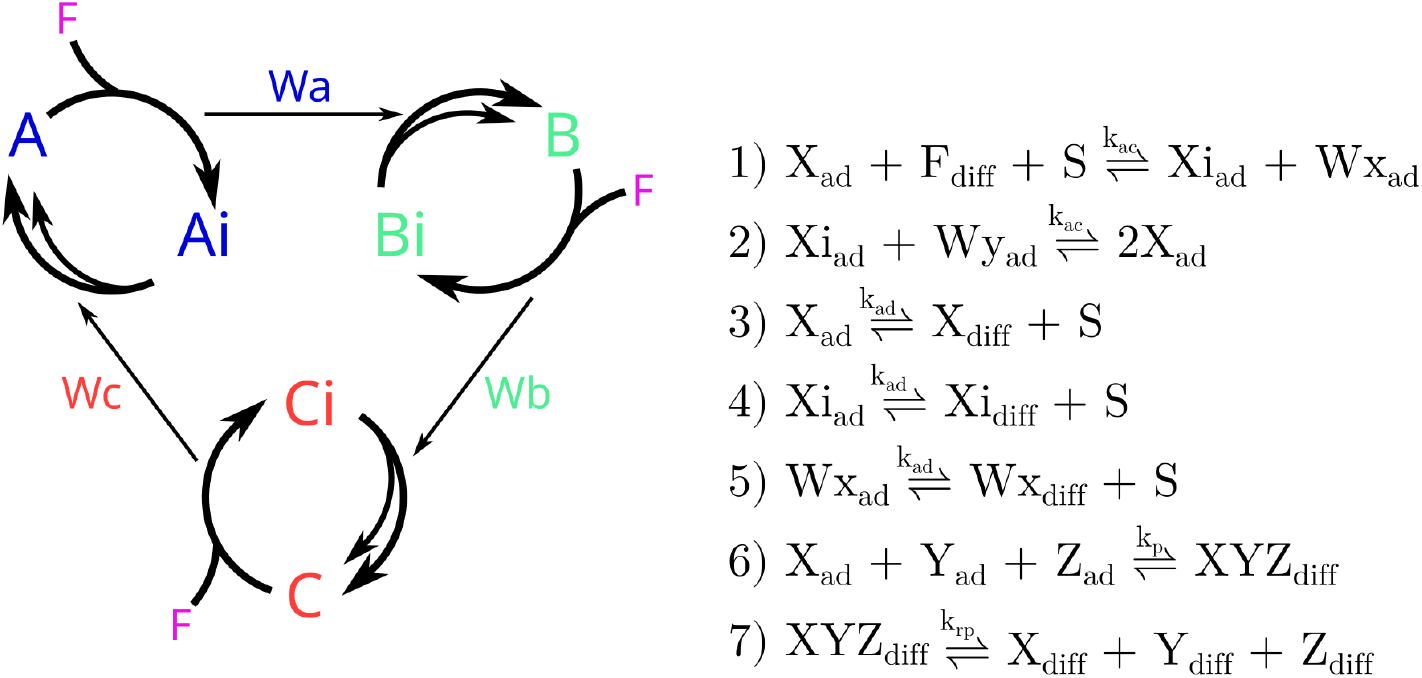
An AC composed of three obligately mutualistic subcycles. Each member, intermediate, and waste molecule can exist in either a reactive adsorbed state (*X*_*ad*_) or a non-reactive desorbed state (*X*_*diff*_). Member species of each subcycle can, when adsorbed, react with a food particle *F* to produce an adsorbed intermediate and an adsorbed waste molecule. The intermediate, when adsorbed, can react with the waste of another subcycle to produce two adsorbed member species. One adsorption site is used up in this process. Reactions 6 and 7, which represent propagule formation and fragmentation, respectively, are only active in cooperatively dispersing cycles. Every species shown is capable of adsorption and desorption except food, which is always found in a desorbed state. In all simulations, *K*_*ac*_ = 0.01, *K*_*ad*_ = 0.01, *K*_*p*_ = 0.01, and *K*_*pr*_ = 0.002. All ACs in our simulations contain three subcycles.

We consider two competing ACs in our models, only one of which forms propagules. We refer to this as the cooperatively dispersing cycle, and to the AC which cannot form propagules as the independently dispersing cycle. These propagules are molecules composed of all the member species of this AC. The two competing ACs are otherwise identical. The propagules detach from the surface as soon as they form, and participate in no reactions aside from dissociation. The reversibility of propagule formation means they can diffuse to new locations and then split to generate the desorbed forms of all the contained member species.

Inspired by Wächtershäuser’s surface metabolism model [15, 16], the ACs in our model are surface-catalyzed, meaning autocatalytic reactions can only occur when the molecules in the cycles are in their adsorbed form. Reversible reactions can transform molecules in the cycles between their adsorbed and desorbed forms, and all molecules except food and propagules can adsorb. Space on mineral surfaces is represented as a finite number of “sites” or *S*. A site is consumed as a reactant in any reaction that increases the number of adsorbed molecules, whereas sites are gained when molecules desorb.

### Stochastic reaction-diffusion model

Our reaction-diffusion model occurs in a two-dimensional world composed of many “pixels” in a hexagonal array with periodic boundaries (Fig 2). The mechanics of diffusion across the array are based on prior models of random walks on hexagonal lattices, as well as methods for coding hexagonal lattices for use in video games [17, 18]. Each pixel functions as a well-mixed reactor in which any allowable reaction can occur. Molecules in their desorbed form can diffuse to one of six neighboring pixels or be lost entirely through outflow reactions, essentially being irreversibly destroyed. The rate of diffusion depends on the mass of the diffusing particle. Since all species but propagules are assumed to have the same mass, propagules will diffuse more slowly relative to the square root of the number of AC member species they contain.

**Fig 2.**
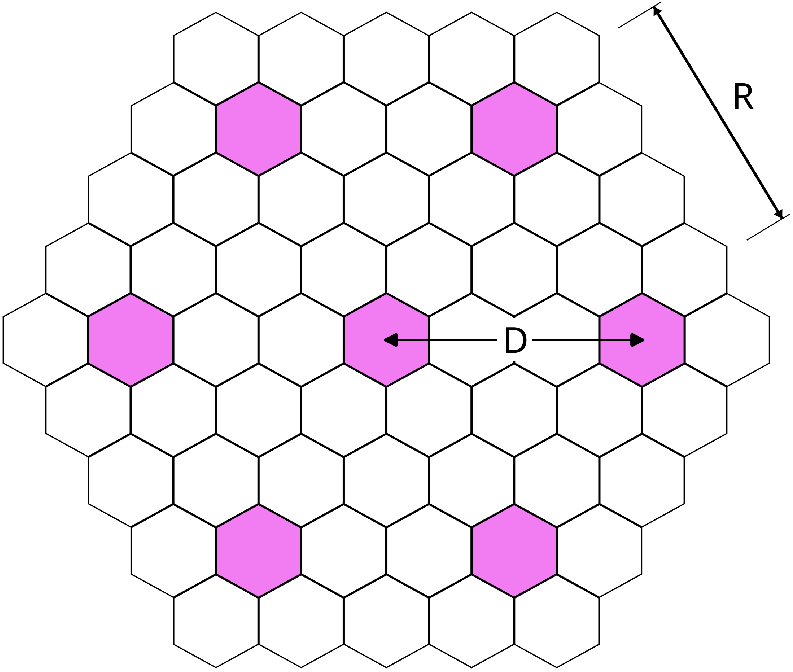
An example hexagonal lattice. The shaded pixels are catalytic mineral patches and the others represent empty space. Each world can be characterized by the distance between catalytic pixels (*D*) and the radius (*R*); here *D* = 2 and *R* = 4. The radius of the world will increase with the value of *D*. The outer boundaries are periodic. In all simulations, the site and food counts in each catalytic pixel are set to *F* = 100 and *S* = 500. Outflow and diffusion occur between neighboring pixels with a rate constant of 0.1, except in the case of propagules, which diffuse and flow out more slowly proportional to their mass with a rate constant of 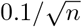, where *n* is the number of subcycles in the cooperatively dispersing AC.

A subset of nineteen evenly-spaced pixels are defined as catalytic sites to which AC member species can reversibly adsorb. By requiring that AC members can only complete the reactions necessary for autocatalysis when in the adsorbed state, we ensure that autocatalytic chemistry is limited to these mineral patches. To allow for density-dependence, which is needed for cycles to gain an advantage by reaching a catalytic site earlier [19], each mineral site contains a limited number of sites *S*. Every catalytic pixel also contains a number of food particles, *F* . *F* is chemostatted in each mineral patch, which amounts to assuming that reaction rates are limited by the availability of surface space but not the availability of food. This aligns with Wächtershäuser’s model, in which carbon dioxide and suitable reductants are abundant and autocatalytic activity is constrained by access to iron-pyrite surfaces [15]. For simplicity, we only model the presence of food in mineral patches and do not track its diffusion throughout the world.

Our model runs using Gillespie’s exact stochastic simulation algorithm [20], which we implemented in MatLab with a tau-leaping approximation [19, 21–23]. Because running third- or higher-order reactions in the Gillespie algorithm can introduce artifacts, we modify the algorithm’s calculation of the rate of the propagule-forming reaction to use the counts of the two reactants with the lowest concentration, rather than all three. Each simulation stops when it reaches a maximum time *T* . We run each simulation long enough to reach an approximate steady state where all the catalytic pixels have been filled.

Though not as fast as using ordinary differential equations (ODEs), the Gillespie algorithm tracks discrete molecule counts. This is important for our needs as it allows for extinction, as opposed to ODEs where concentrations asymptotically approach zero. Moreover, discrete counts are essential here as our study system involves ACs whose dispersal ability is limited by the presence or absence of small counts of member species, and ODEs can produce misleading results in systems where molecule counts are very low [24].

### Disturbances

Some simulations also included environmental disturbances, modeled as random events in which a single catalytic pixel is wiped of all its contents, leaving nothing but the original number of sites *S* and food *F* . A physical scenario yielding such an outcome would be the surface of a mineral patch flaking off or being washed clean, leaving behind a fresh surface of the same size. Disturbances occur at a frequency based on an exponential distribution with a mean *λ*, which is the average time between disturbance events.

### Relative concentrations and extinction rates

In simulations where cooperatively dispersing cycles are competing with independently dispersing cycles, we assess their success by calculating their relative concentration. The relative concentration, *A*_*rc*_, is the total count (across the entire world) of all the member species molecules of the cooperatively dispersing cycles divided by the total count of all member species molecules of both cycles. An *A*_*rc*_ of 0.5 means both cycles are equivalent in abundance whereas higher values mean the cooperatively dispersing cycle is more prevalent in the world. We also record whether one or both cycles goes globally extinct during a simulation.

## Results

### Well-mixed environments

Propagule formation has a cost because it siphons away member species that would otherwise undergo autocatalytic reactions. Therefore, in a well-mixed environment, if all reaction rates and starting concentrations are held equal, a cooperatively dispersing cycle should be at a disadvantage relative to one that does not form propagules. We verified this intuition by running simulations seeded with equal concentrations of the two ACs in a world with only one catalytic pixel. As anticipated, when the rate of propagule formation is higher than zero, cooperatively dispersing cycles tend to fall to a lower relative steady-state concentration. As the rate of propagule formation increases, this steady-state concentration approaches zero (Fig 3).

**Fig 3.**
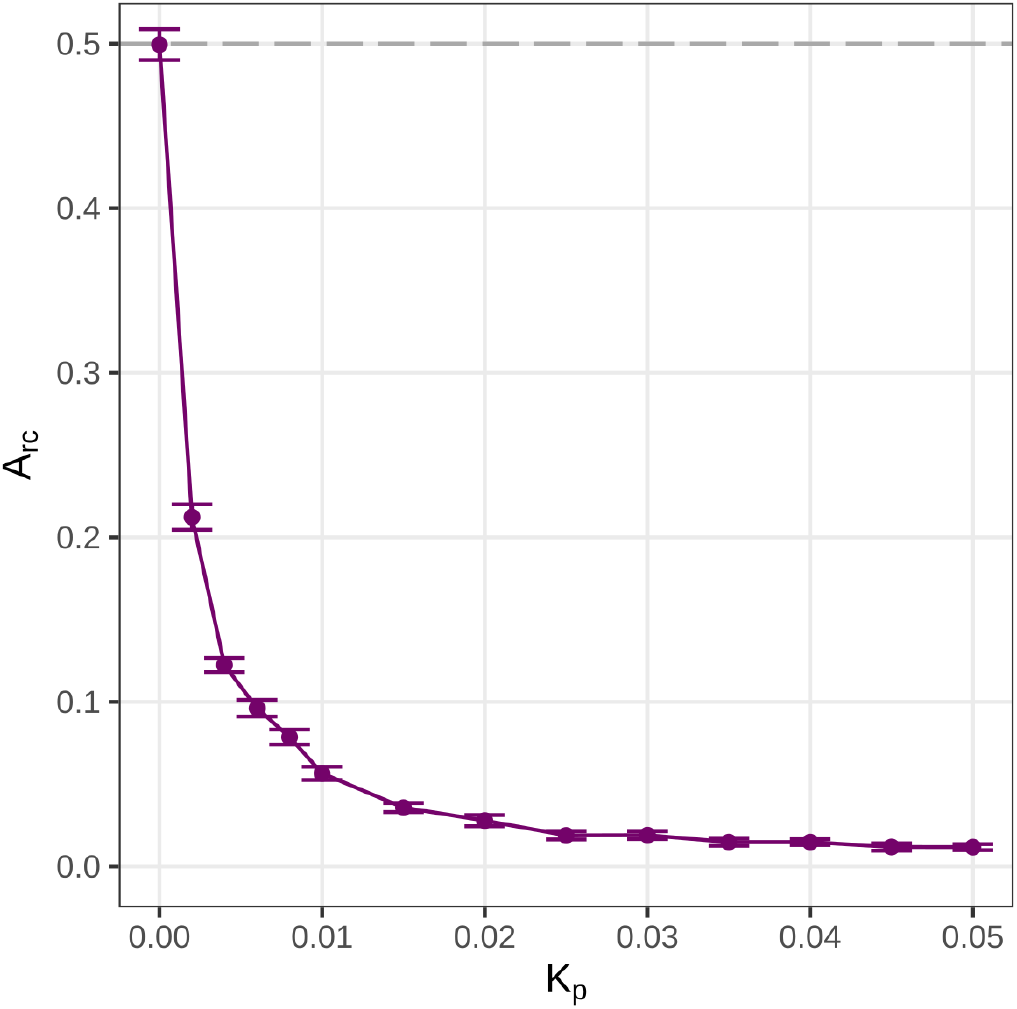
Relative concentration (*A*_*rc*_) of the cooperatively dispersing cycle with different propagule formation rates (*k*_*p*_) in a well-mixed world with no spatial structure. When *k*_*p*_ = 0 there is on average no difference between the concentrations of the two cycles; as *k*_*p*_ increases *A*_*rc*_ approaches zero. Error bars represent standard error with 25 replicates per point.

### Spatial environments

We then ran simulations in worlds with multiple catalytic sites separated by empty space. Every catalytic pixel is separated from its nearest neighboring catalytic pixels by a distance *D*, which is the minimum number of dispersal steps needed for a particle to travel from one to another. Simulations were initiated with one cooperatively and one independently dispersing cycle seeded in two random catalytic pixels. Each cycle started with 25 of each of their member species in an adsorbed state. The remaining catalytic pixels contained only empty sites and food. For each value of *D*, we ran each simulation until it reached a steady state, defined as a state in which approximately all catalytic sites have been filled and there are no longer any clear changes in the concentration of any species beyond fluctuations around a mean. The necessary lengths of time were found through trial and error in preliminary simulations and increased with the distance between catalytic sites.

We find that, as *D* increases, the final relative concentration of the cooperatively dispersing cycle also increases, surpassing that of the independently dispersing cycle when *D* = 3 (Fig 5). Notably, neither cycle ever drives the other extinct. However, while both cycles coexist in the world, they tend not to coexist within mineral pixels (Fig 4). The lack of coexistence within pixels arises because high concentrations of member species, intermediates, and waste of a cycle can effectively block the adsorption of the opposing cycle to a mineral patch. Such bistability, manifesting here as a priority effect, was previously reported in a similar model of surface-limited autocatalysis [19]. The pattern we observe displays a degree of robustness to changes in experimental parameters; we reach similar results in simulations composed of ACs with 2 or 4 subcycles (S1 Fig), as well as those where interactions with subcycles are facultative rather than obligately mutualistic (S2 Fig).

**Fig 4.**
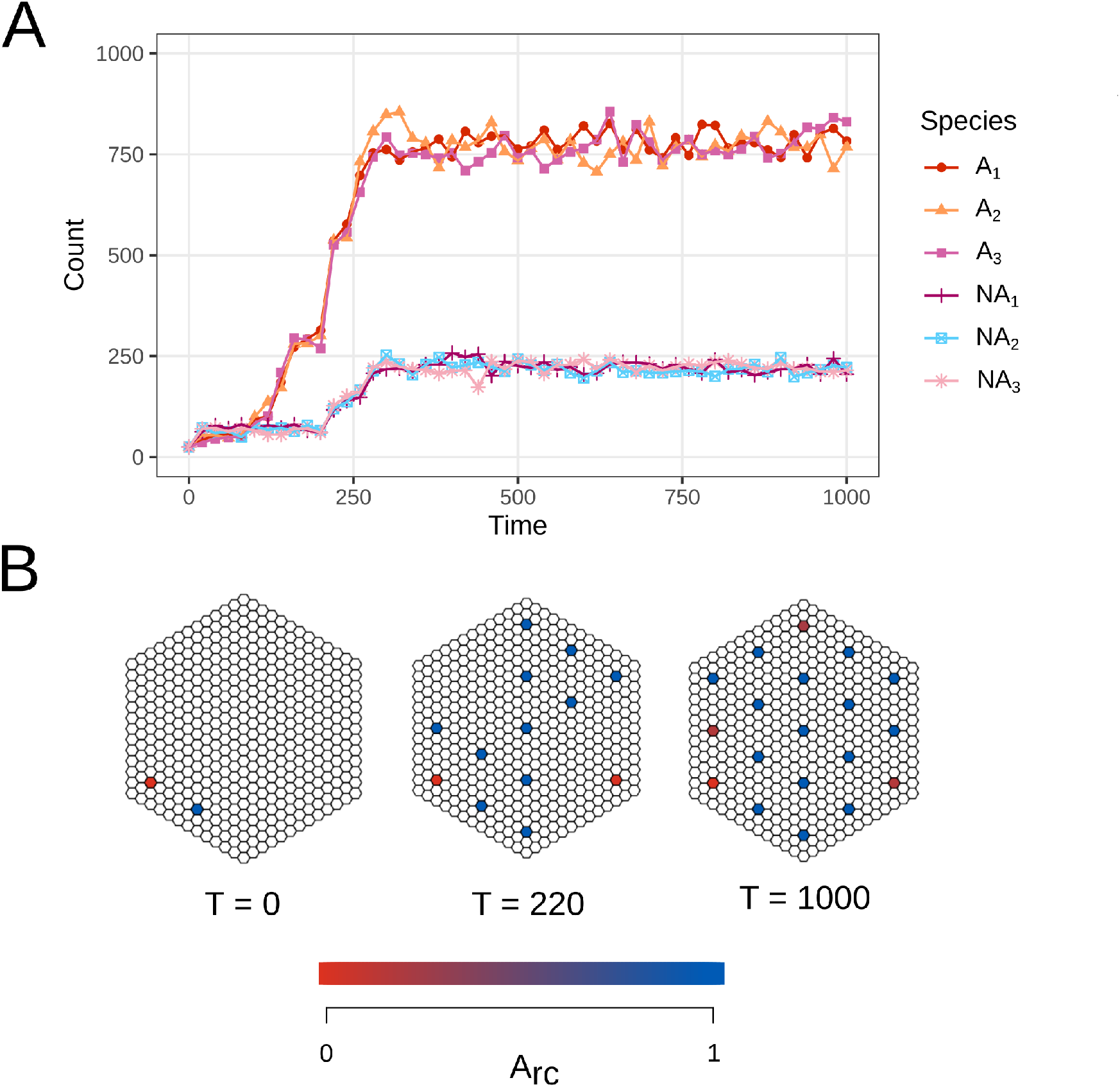
Example run where *D* = 5. As the simulation runs, the global counts of the member species of each cycle tend to remain stable until a new catalytic pixel is seeded, at which point the counts rapidly increase until the carrying capacity *S* of the newly seeded pixel is reached (A). Once all the sites have been filled, the global counts of each cycle fluctuate but remain approximately stable. In this example, the cooperatively dispersing cycle (species *A*_1_ - *A*_3_) seeds more catalytic seeds than its independently dispersing competitor (species *NA*_1_ - *NA*_3_), quickly dispersing throughout the world. As visualized in (B), there is little coexistence between the cycles within pixels; instead, they tend to become completely and permanently filled by whichever cycle first seeded them.

**Fig 5.**
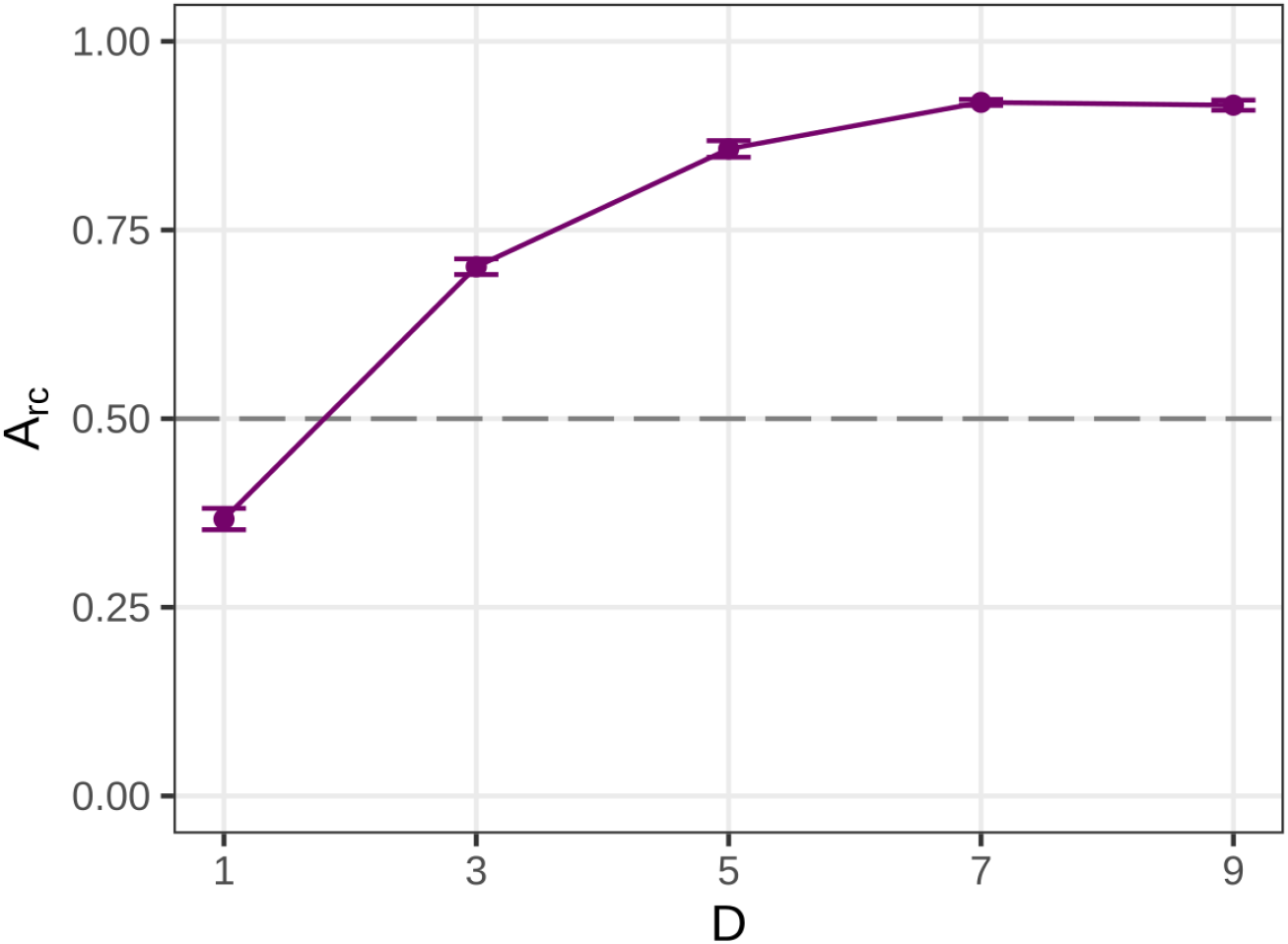
Relative concentration of the cooperatively dispersing cycle based on the distance between each site. When D = 1 and sites are directly adjacent to each other, propagule formation is disadvantageous and results in low relative concentrations. However, cooperatively dispersing cycles reach higher relative concentrations as D increases, leveling off at approximately D = 7.

### Disturbance

We next examined the effect of adding environmental disturbances. We predicted that, in the presence of disturbance, propagule formation would act as a mechanism of ecological resilience, allowing the cooperatively dispersing cycles to escape catalytic sites before being locally extirpated. We ran simulations for logarithmically increasing values of the disturbance frequency *λ*, ranging from 10 to 10000. Simulations were run for 9000 time steps, meaning that for the lowest value of *λ* disturbances were extremely frequent and for the highest frequency there might only have been a single disturbance event, if any, in the entire run.

For higher values of *D*, the advantage of propagule formation is more pronounced than in simulations with no disturbance (Fig 6). Additionally, extinction becomes more common as disturbances become more frequent (Fig 7). Generally, the independently dispersing cycle is more likely to go extinct, though the cooperatively dispersing cycle is also vulnerable to extinction at low values of *D*.

**Fig 6.**
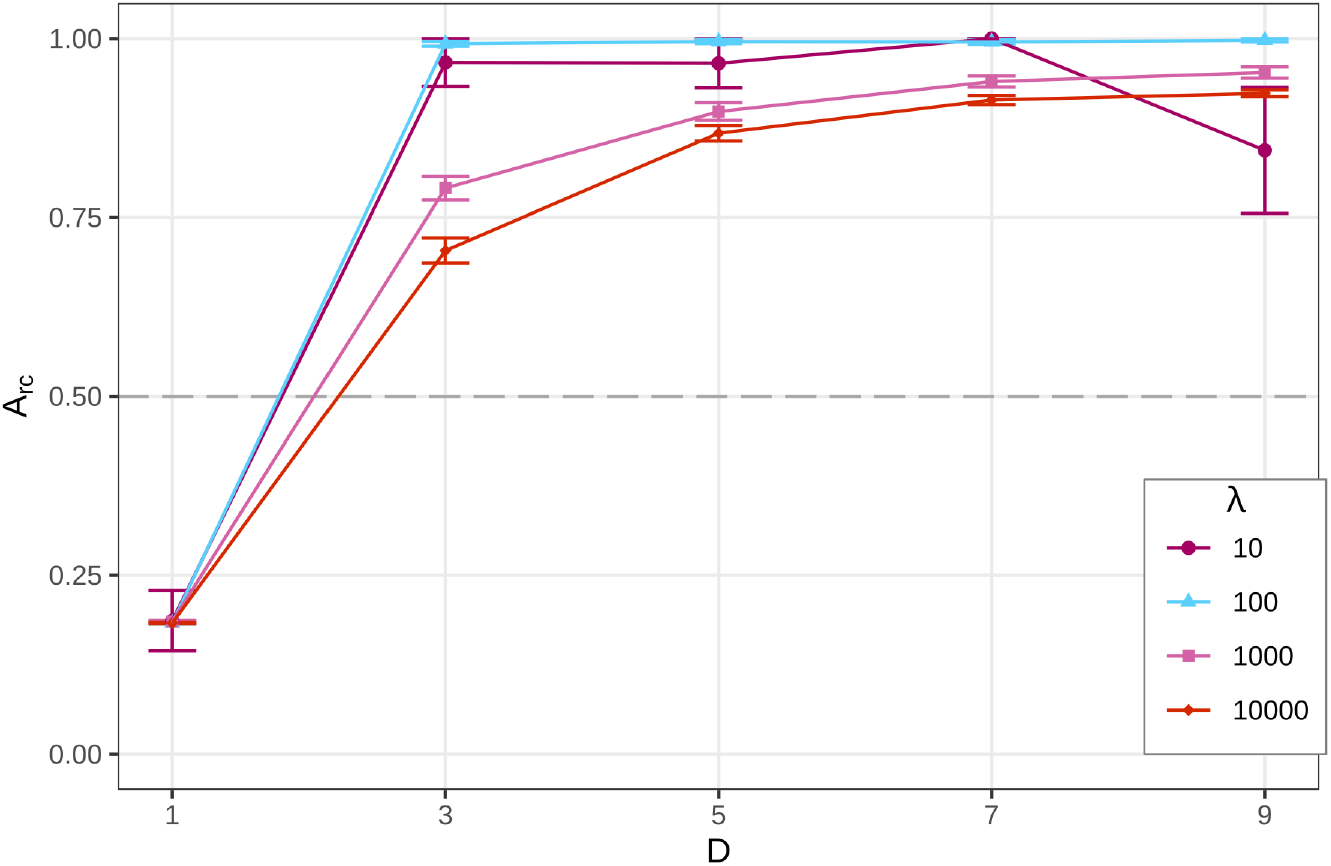
Relative concentration of the cooperatively dispersing cycle based on disturbance frequency. Disturbances are less frequent as lambda increases. As in the undisturbed simulations, *A*_*rc*_ rises as the distance D increases, though it tends to reach higher values when disturbance is more frequent. Simulations where both cycles go extinct are not included in the calculation of *A*_*rc*_.

**Fig 7.**
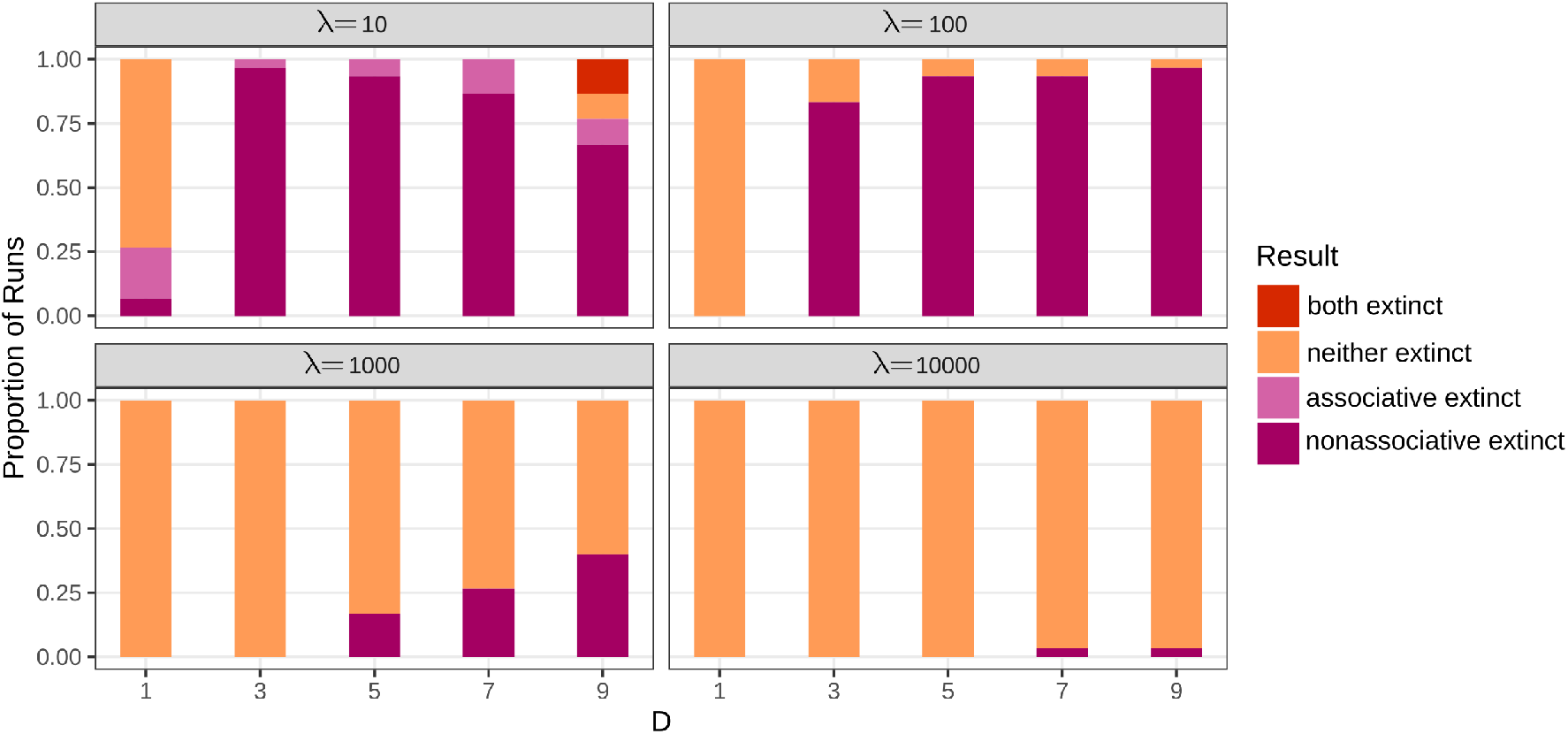
Extinction rates for each combination of D and disturbance frequency. At lower disturbance frequencies, extinction is rare, but becomes more likely as the disturbance frequency increases. In cases where extinction occurs, the cooperatively dispersing cycle is much less likely to go extinct, except when *D* = 1.

## Discussion

Our results demonstrate that spatial structure and disturbance change the relative success of competing autocatalytic motifs and support chemistries that would be unlikely to persist under well-mixed conditions.

In this way it reaffirms prior work on spatially supported chemistries and selection upon autocatalytic cycles [19].

Codispersal is especially advantageous when catalytic sites are widely separated and environmental disturbances are frequent. This pattern supports the cells-as-propagules hypothesis, which proposes that the first evolving systems were autocatalytic reaction networks adhered to mineral surfaces with protocells deriving from propagules [11]. In patchy, extinction prone environments, cycles that are better at evading extinction and dispersing would be expected to dominate over time [19]. Our data show that, when cycles are composed of cooperating subcycles, these attributes can be conferred by the production of simple propagules. The tendency for propagule formers to become more abundant over time resembles selection, but since it lacks either autopoiesis or genetic mechanisms it blurs the line between paradigmatic evolution and ecosystem dynamics.

Though our model represents surface-catalyzed small-molecule autocatalysis, this is probably not the only condition under which codispersing systems would tend to become enriched. For instance, nucleic acids exhibit codispersal insofar as duplexes carry both complementary strands needed for replication. Moreover, a long nucleic acid strand may carry many distinct genes, effectively being a propagule for multiple separate functional sequences. It is also worth noting that some RNA-based prebiotic scenarios rely on hypercycles of cooperating ribozymes [25–27], which might also drive the enrichment of codispersal mechanisms. In general, we expect any mutualistic set of replicators, placed in a spatially structured environment with distantly spaced resources, to have a tendency to acquire the ability to cooperatively disperse.

Our work also has implications for the role of side reactions in prebiotic chemistry. Side reactions are sometimes considered a major obstacle to the viability of pre-enzymatic autocatalysis, as they siphon molecules away from autocatalytic cycles [28]. Propagule formation is a side reaction in our system. Nonetheless, despite being a drain on propagule-forming cycles in a well-mixed system, it can be beneficial when a spatial structure with sparsely distributed resources is introduced. This aligns with other analyses showing that side reactions may create reservoirs from which autocatalysts can be replenished and have diverse effects on autocatalytic systems [29]. Our results also demonstrate the importance of considering spatial structure when modeling prebiotic chemistry; networks that seem unfavorable for the stability of autocatalytic cycles in well-mixed environments may show different behavior when a spatial structure is present [30].

By analyzing the effect of spatial structure and disturbance on enrichment for codispersal ability, our model lays a groundwork for further work on the emergence of protocells from propagules. It is important that future work consider larger lattice sizes and more complex (and realistic) reaction networks to ensure that the principles explored here hold under chemically realistic conditions and when ecological dynamics other than simple mutualism and competition exist. Additionally, much work remains to demonstrate that selection for codispersal can result in propagules with the capacity to evolve into protocells.

Important next steps are to test the behavior of different types of propagules such as vesicles or coacervates. It is then vital to demonstrate that propagules themselves can undergo selection. Similar to the evolution of multicellularity or endosymbiosis, the emergence of propagules as nascent individuals may result in two conflicting levels of selection [31] – on the surface-adsorbed ACs, and the bounded structures they cooperatively produce. If this happens, selection for effective and more tightly integrated propagules would need to outweigh selection for faster-replicating or otherwise better-persisting autocatalytic subcycles. This shift in levels of selection, though completely abiotic, would be the first major transition in individuality.

## Supporting information

**Fig S1.**
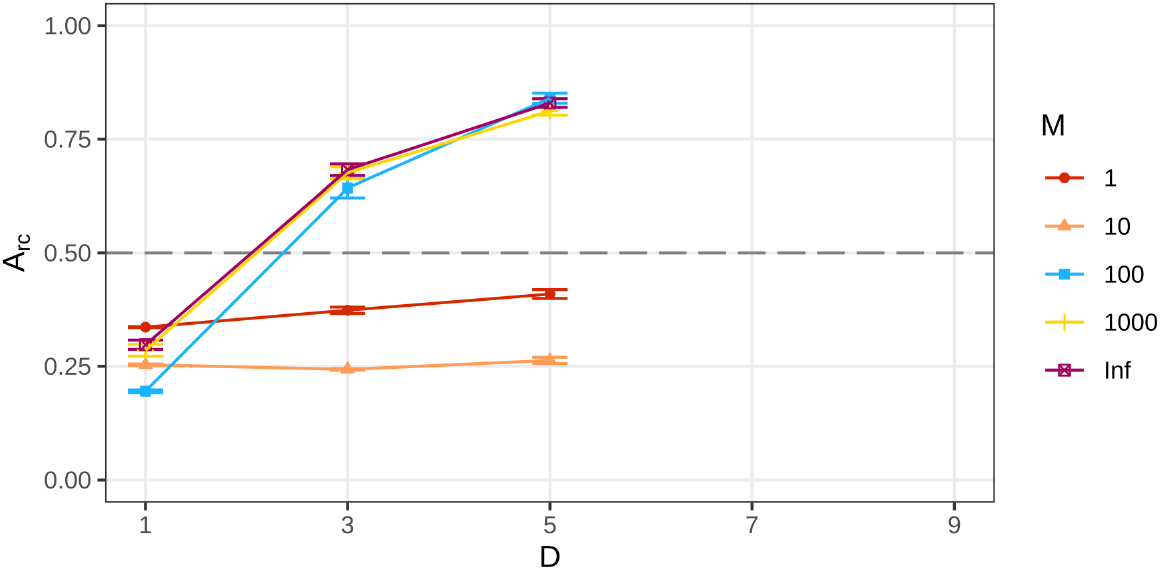
*A*_*rc*_ when subcycle mutualism is facultative. The interdependence of subcycles can be relaxed by adding an extra reversible pathway for the synthesis of a member species from an intermediate, which requires a generic food molecule *F* rather than the waste of another cycle. The rate constant of this reaction is controlled by a parameter *M*, which is the number of times slower this reaction is relative to the reaction that uses the waste of another subcycle (*K*_*ac*_). Propagule formation is disadvantageous, and the *A*_*rc*_ curves flatter, when the generic pathways are roughly as fast as the mutualistic pathway. When generic pathways are slower, the curves converge on those of the obligate case. Simulations are only run to *D* = 5 here as simulations with longer distances become computationally intractable with the additional reactions that need to be tracked.

**Fig S2.**
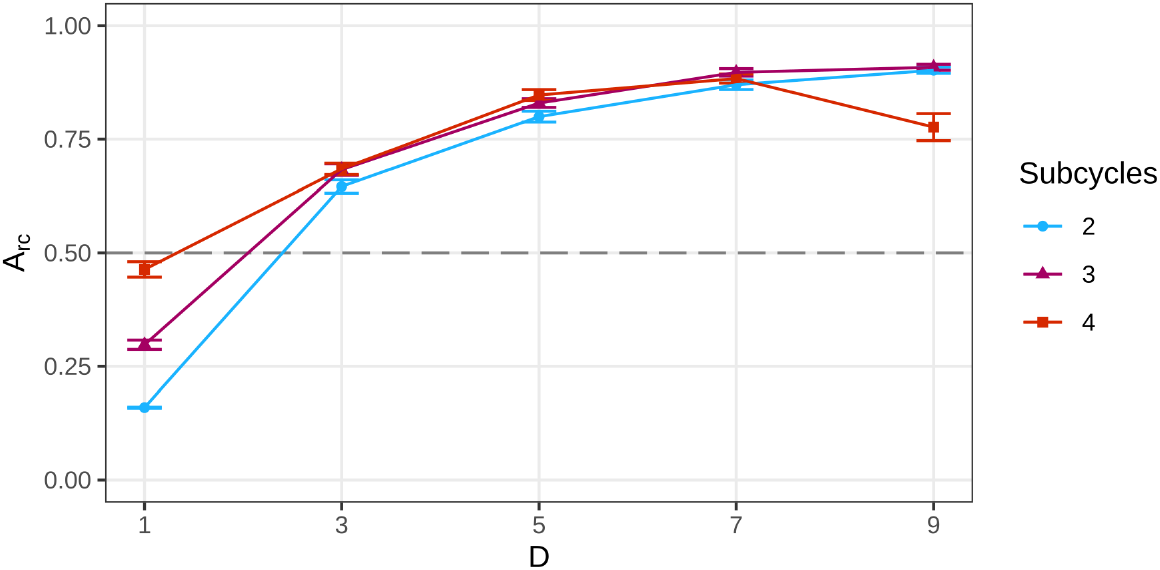
*A*_*rc*_ for systems with 2 or 4 subcycles. *A*_*rc*_ curves are qualitatively similar regardless of whether ACs contain 2, 3, or 4 subcycles. At short distances, the relative advantage of the cooperatively dispersing cycle over the independently dispersing cycle appears to increase only slightly as the number of subcycles increases, and is most pronounced at *D* = 2. Additionally, the effectiveness of propagule formation in the most complex cooperatively dispersing cycle appears to collapse at *D* = 9. Further work is needed to fully characterize how cycle complexity influences codispersal dynamics in competitive systems, especially when cooperatively dispersing and independently dispersing cycles might contain different numbers of subcycles.

## Acknowledgments

The authors would like to thank Tony Ives, Betül Kaçar, Zoe Todd, and John Yin for feedback on this study. We also thank Alex Plum, Zhen Peng, Tymofii Sokolskyi, Emily Spencer, and A. Khanov for discussions about prebiotic chemistry in spatial environments. Computation was made possible by the Center for High-Throughput Computing at the University of Wisconsin-Madison [32]. Baum acknowledges support from the National Science Foundation under Grant No. 2218817. This material is also based upon work supported by the National Science Foundation Graduate Research Fellowship Program under Grant No. 2137424. Any opinions, findings, and conclusions or recommendations expressed in this material are those of the authors and do not necessarily reflect the views of the National Science Foundation.

## Code availability

Code and data used in this paper are available at https://github.com/NA-ori/molecular_codispersal.

## Notes

### Competing Interest Statement

The authors have declared no competing interest.

